# Integrating indigenous knowledge, ontology, and molecular barcoding to characterize spider monkey (*Ateles paniscus*) filariasis

**DOI:** 10.1101/2020.10.26.354985

**Authors:** Marissa S. Milstein, Christopher A. Shaffer, Laramie L. Lindsey, Tiffany M. Wolf, Philip Suse, Elisha Marawanaru, Evan J. Kipp, Tyler Garwood, Dominic A. Travis, Karen A. Terio, Peter A. Larsen

## Abstract

Zoonotic disease risk is greatly influenced by cultural practices and belief systems. Yet, few studies have integrated traditional ecological knowledge and ontology with western ways of knowing to better understand potential zoonoses. Here, we integrate molecular phylogenetics, histopathology, and ethnography to characterize a filarial nematode found within the abdominal cavity of spider monkeys (*Ateles paniscus)*. The filarid is recognized as ‘spider monkey cotton’ by indigenous Waiwai in the Konashen Community Owned Conservation Area, Guyana. Ethnographic data revealed that the Waiwai perceive of ‘spider monkey cotton’ as a normal characteristic within the ‘spider monkey person.’ Further, the Waiwai indicated that ‘cotton’ was ubiquitous with spider monkeys and is not understood to be infectious nor zoonotic. This distinction is in contrast to other internal parasites found within spider monkeys that the Waiwai know to cause disease in both monkeys and humans. Our morphological and molecular characterization support the classification of the filarid as *Dipetalonema caudispina*, a minimally studied and seemingly non-zoonotic parasite, consistent with Waiwai beliefs. Thus, our findings allow us to establish commensurability between scientific knowledge and indigenous ontology. More broadly, this work highlights the importance of integrating multiple knowledge systems and leveraging advanced genomics to better understand and prevent emerging zoonotic diseases.

## Introduction

The hunting of wildlife is critical to the food security and livelihoods of millions of people throughout the tropical world [1,2]. This is particularly true in Amazonia, where wild meat is a major source of protein [3–5], fat [5], and micronutrients [6]. Beyond economic importance, subsistence hunting plays an integral role in the social and symbolic lives of millions of Amazonians [7–9]. Unfortunately, the hunting of wildlife also presents an especially high risk interface for the emergence of zoonotic disease. The frequent contact between hunters and wildlife, and the handling of bodily fluids during butchery and consumption, greatly increases the potential for pathogens to cross the species boundary [10,11]. Importantly, this interface, and the potential risk for zoonotic transmission, is highly influenced by the cultural practices and beliefs surrounding wild meat hunting [12].

Over the past few decades, considerable research has been dedicated to identifying potential zoonoses from wild meat and characterizing practices that may increase the risk of pathogen transmission [11,13–16]. Collectively, these studies have used a variety of methods to identify likely zoonoses and assess the risk of zoonotic disease transmission, with methods including: ecological modeling, molecular screening of wild meat samples, and community survey-based approaches. However, few studies have sought to incorporate the ecological knowledge of local or indigenous populations to characterize zoonotic disease risk from wild meat [17]. Local ecological knowledge is, “the knowledge that people in a given community have developed over time, and continue to develop. It is based on experience; often tested over centuries of use; adapted to the local culture and environment; embedded in community practices, institutions, relationships and rituals; held by individuals or communities; and dynamic and changing” [18]. The concept of local ecological knowledge is somewhat more broad than the closely related concepts of traditional ecological knowledge (TEK) or indigenous ecological knowledge, as it includes knowledge of groups that may not qualify as “indigenous” and avoids the difficulties involved in determining when knowledge is “traditional,” (17). Western scientists have increasingly recognized the importance of local ecological knowledge for documenting pharmaceutical benefits of plants [19], monitoring and adapting to environmental change [20], and conserving biodiversity [21]. Several recent studies have shown the value of local ecological knowledge for understanding and combating emerging infectious diseases [17,22,23]. Yet, local ecological knowledge has rarely been incorporated in studies of zoonotic risk from wild meat.

Similarly, few researchers have incorporated ontology, the way a society constructs reality, in studies of zoonotic disease [24,25]. As the nature of being and reality, ontology is related to but distinct from knowledge. Studies of local ecological knowledge generally focus on knowledge that is easily placed within western constructions of reality (e.g. knowledge of animal movement, climate change, medicinal qualities of plants) rather than seeking to also understand how these societies conceptualize being and reality when their knowledge systems (and not just the components of knowledge) differ from Western metaphysics [24,25]. For example, it is much easier for scientists to integrate scientific and indigenous knowledge on jaguar ecology when they both agree that a jaguar is an animal and not a person. However, finding commensurability becomes more difficult if one ontological system identifies both as persons - with jaguars having similar cultures, social practices, and rituals as humans, plants, water, and other persons - while the other system makes clear biological and cultural demarcations between jaguar and human [17].

Understanding how people who are highly dependent on wildlife for food conceptualize human-animal-environment relationships and make meaning from their worlds may provide important information about potential zoonotic risk. Societies that have occupied the same environment for long periods of time may have developed belief systems that act to prevent the spread of zoonotic pathogens. In addition to providing information that may help mitigate the emergence of zoonoses, characterizing ontological divergence can also demonstrate how the different ontologies of groups living in different environmental and cultural contexts can be integral to their long-term health and social and political self-determination. According to Ludwig and colleagues, “Divergent ontologies therefore do not indicate that indigenous communities fail to grasp the structure of the natural realm but rather reflect how different cultural and explanatory practices come with different ontological requirements” [25](p.43).

Many indigenous societies hold that all animals are persons, with similar agency and sense of self as humans, but with different bodies [26–30]. In the belief systems of these “perspectival multinaturalist” societies, all beings see themselves as persons just like humans see themselves as persons [28]. However, distinct beings (e.g. a jaguar, a human, a monkey) see each other as distinct because of their different external bodies. Morphological and physiological traits of different species are believed to represent species specific “clothes” that mask the underlying similarities that unite all beings as persons. As these societies attribute cultural elements similar to their own to animal persons, they often view hunting as a generalized reciprocity with the animal world, with animal prey being offered as gifts and continued prey availability governed by these gifts being properly received [31–34]. Thus, wild meat “gifts” must not be refused (i.e. animals should be hunted when they are available) and wild meat hunting represents an integral social, as well as economic practice. Understanding how the worldviews of these societies differ from a Western metaphysical worldview is essential for mitigating disease risk and informing potential health interventions. Yet, in many ways, this view of the connectedness among all animals and the porosity of species boundaries is highly consistent with the scientific understanding of zoonotic disease and the One Health framework.

One society that holds perspective multinaturalist beliefs are the Waiwai, a group of forager-horticulturalists in Southern Guyana and Northern Brazil [35–38]. In 2016, MSM and CAS, in collaboration with the Waiwai community, initiated a hunter-based surveillance program for mitigating zoonotic disease from wild meat. During the initial stages of this program, MSM and CAS observed extensive filarid infections in the peritoneal cavities of all spider monkeys (Fig 1a). These filarids were described by the Waiwai as ‘*poroto maúre* or ‘spider monkey cotton’, a normal characteristic of spider monkeys and that spider monkeys use the ‘cotton’ to spin into *‘keweyu’* or aprons, a traditional garment worn by Waiwai women (Fig 1b). However, some Waiwai hunters also asked the researchers (MSM and CAS) what they thought the ‘spider monkey cotton consisted of and whether they thought it could be passed to humans. Therefore, we sought to identify the filarid while working within the belief system of the Waiwai by acknowledging the connection between the filarids and the spider monkey person.

**Fig 1.**
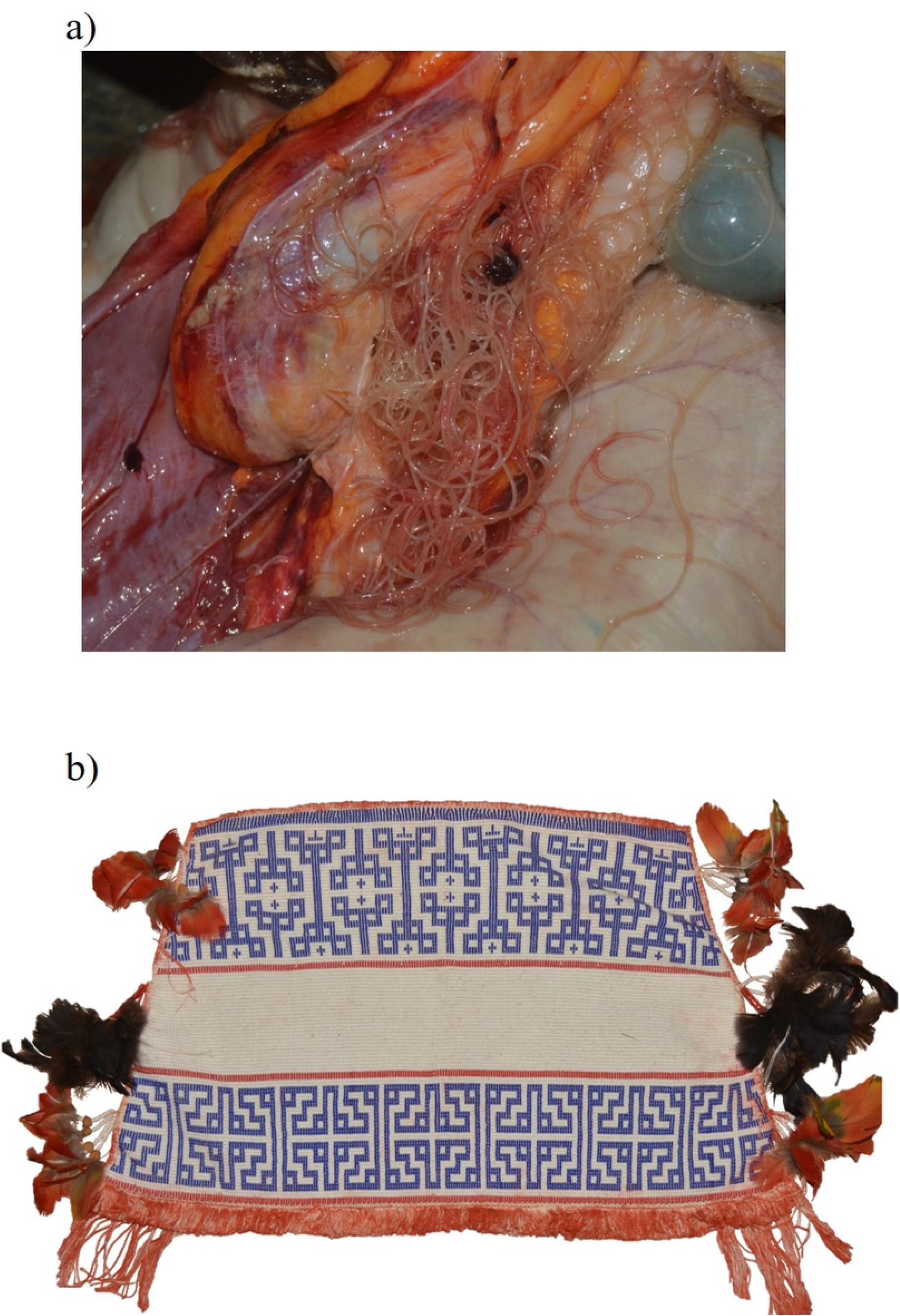
a) Filarial infection within the peritoneal cavity of a black spider monkey (*Ateles paniscus*). The Waiwai refer to the filarids as *poroto maúre* or ‘spider monkey cotton’ and state that the ‘cotton’ is used by the spider monkeys to construct aprons (kewey*ú*). **b) Traditional Waiwai apron**. Apron made from hand-woven cotton and traditionally worn by women.

The overarching goal of this study was to characterize the filarial nematodes identified by the Waiwai as spider monkey cotton from both a western evolutionary perspective and from the ontological understanding of the Waiwai. Our mixed-methods study had two primary objectives; 1) to characterize Waiwai understanding of spider monkey cotton, its pathogenicity and zoonotic potential through ethnographic engagement and 2) to determine taxonomic placement of the filarial nematode through histopathology and both traditional Sanger sequencing and newly developed nanopore sequencing technology (i.e., Oxford Nanopore Technologies MinION Flongle sequencing). Using this integrated approach, we sought to better understand a culturally significant connection between the Waiwai and the Amazonian fauna and to assess commensurability between divergent knowledges and ontologies. Further, employing nanopore sequencing technology alongside ethnographic data collection allowed us to assess the practicality of future deployment of a mobile sequencing lab to Southern Guyana for real-time molecular barcoding of micro- and macro-parasites.

## Material and Methods

### Study site

The Konashen Community Owned Conservation Area (KCOCA) is a 625,000 ha indigenous reserve in southern Guyana, South America owned and managed by indigenous Waiwai forager-horticulturalists (Fig 2). Approximately 225 Waiwai live within the KCOCA, concentrated in the village of Masakenari. The Waiwai subsistence strategy consists of slash and burn cassava horticulture supplemented by a high dependence on wild meat from a wide variety of species [9,39,40]. Primates are among the most frequently consumed animals for the Waiwai, particularly the black spider monkey (*Ateles paniscus*) [9]. Current Waiwai beliefs were highly influenced by missionaries from the unevangelized field mission in the mid 20th century and most Waiwai now strongly identify as Christian. However, in concert with their continued reliance on traditional subsistence strategies and intimate connection with wildlife and the natural world, the Waiwai retain many aspects of the perspective multinaturalist ontology [9].

**Fig 2.**
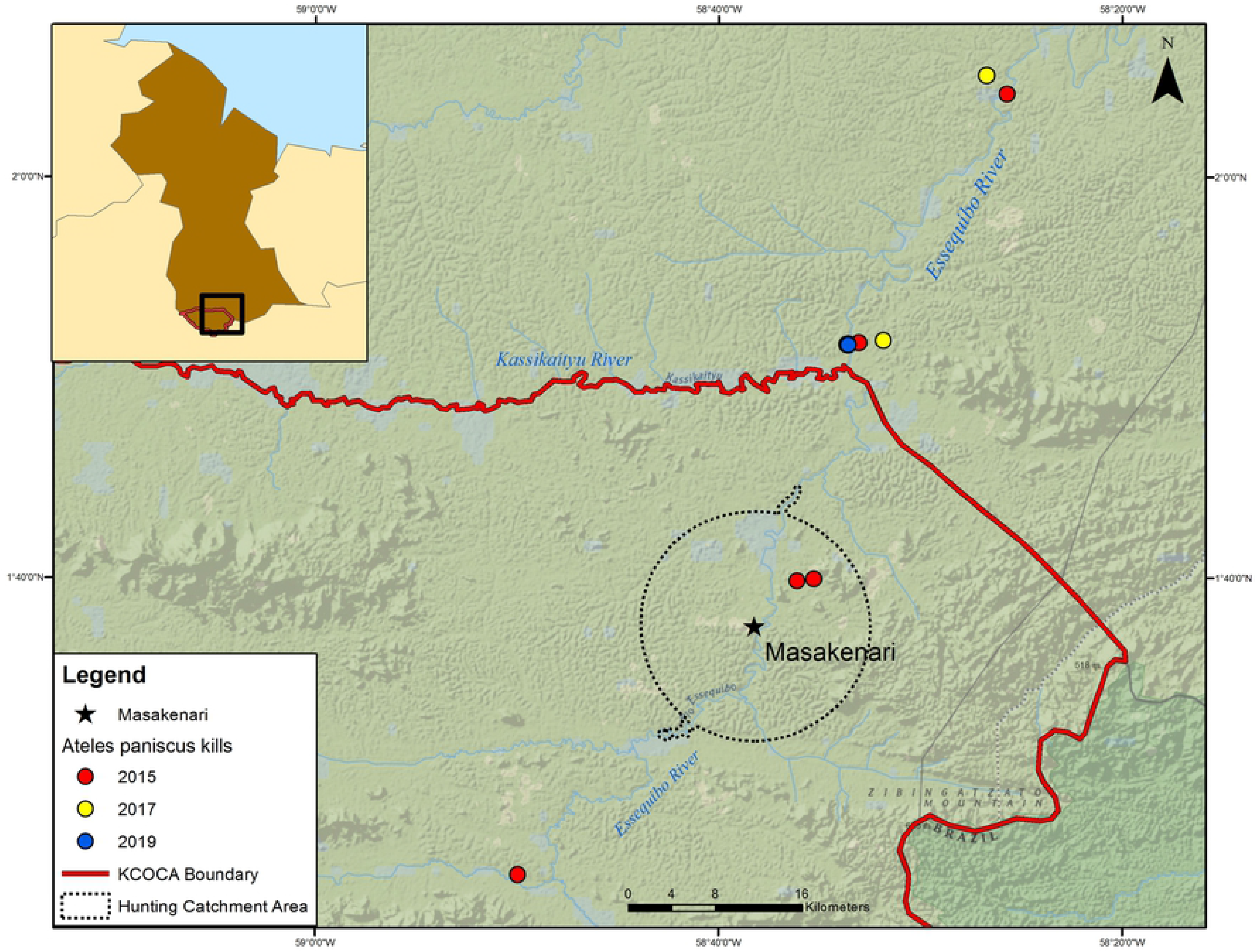
Konashen Community Owned Conservation Area (KCOCA). Boundary of the KCOCA outlined in red. Waiwai village of Masakenari depicted by the star. Waiwai hunting catchment area outlined by the dotted line. Location of each spider monkey harvested by Waiwai hunters included in the analysis depicted by red (2015; n =5), yellow (2017; n =2) and blue (2019; n =4) dots. All samples from 2019 were collected in the same location.

### Ethnographic Methods

We collected qualitative data as a part of an ongoing ethnographic study with the Waiwai during three study periods from June-August 2015, July-August 2016, and June-August 2017 [9,12]. To assess Waiwai perceptions of the filarial infections and elucidate their beliefs about the pathogenicity and transmissibility of the parasites, we used a combination of semi-structured and unstructured interviews, and participant observation. Interviews were conducted in either Waiwai, English, or a combination of both. In cases where the interviewee was not fluent in English, a Waiwai field assistant acted as an interpreter.

We conducted semi-structured interviews with 30 individuals as part of a concurrent study of zoonotic disease risk [12]. Participants were recruited through snowball sampling where the Waiwai Village Council initially suggested five hunters to interview and these hunters provided names of five more hunters for a total sample of 30 hunters. During these interviews, we asked participants about the occasions when they observed filarid infections (i.e. spider monkey cotton) and which animal species these infections were associated with. We conducted unstructured interviews opportunistically during participant observation of hunting and butchery activities. Each time we observed filarids in the peritoneum of spider monkeys (n = 47), we asked those involved in the butchery to identify the filarids and describe how the animals acquire the parasites.

We also conducted in-depth unstructured interviews with four key informants (two male and two female) on the broader topic of etiology of disease and beliefs about wildlife [9,12]. We identified informants by initially asking the Waiwai Village Council for suggestions about potential informants. We then began conducting interviews and participant observation with these individuals to ascertain the extent to which they would be suitable informants. Semi-structured and unstructured interviews were transcribed in English from field notes each day during field seasons. We used grounded theory to extract meaningful themes and repeated elements from semi-structured and unstructured interviews and applied open coding to our data [41]. Themes were identified by assessing the frequency, context, co-occurrence, and other patterns of concepts. Coding was performed by CAS.

### Sample collection

Adult filarial worms were collected from the intraperitoneal cavity of spider monkeys shot by Waiwai hunters during two, three-month sampling periods between June to August 2015, 2017, and a two week sampling period in July 2019. Samples were collected during normal butchery activities by Waiwai hunters and no spider monkeys were killed specifically for this research. For each spider monkey butchered, a sample of the intraperitoneal nematode population was collected (2015, n=5; 2017, n=2; 2019, n=4) and stored in RNA-later. All samples were either maintained at room temperature (2015, 2017) or liquid nitrogen (2019) until transferred to a −80C freezer at the University of Minnesota College of Veterinary Medicine.

### Histopathology methods

Representative samples of liver, kidney, heart, lung and spleen were collected and fixed in either RNAlater or 10% neutral buffered formalin for histopathology from the same spider monkeys from which nematodes were collected in 2015 and 2017 (samples collected in 2019 were not subject to histopathology). Sections were routinely processed, embedded in paraffin, and 3-5 micron sections were stained with hematoxylin and eosin for histologic evaluation.

## Molecular methods

### DNA extraction, PCR, and sequencing

DNA was extracted from subsamples of nematodes (approximately 2 to 3 nematodes per subsample) collected from 11 individual spider monkeys using a DNeasy commercial kit following the manufacturer’s instructions (Qiagen, Germany). Prior to extraction, approximately 25mg of nematode tissue was treated with 180ul of Buffer ATL, 20ul of Proteinase K and 2ul of a 5% DX solution for two, 30 second cycles at 2500rpm with a 30 second pause between cycles using a TissueLyser II (Qiagen, Germany). Samples were then incubated at 56C for 15 to 60 minutes until completely lysed and stored at 4C until use.

Molecular barcoding focused on the mitochondrial cytochrome c oxidase subunit I (*CoxI*) gene using the following PCR primer sequences from Lefoulon and colleagues: FCo1extdF1 (forward): TAT-AAT-TCT-GTT-YTD-ACT-A and FCo1extdR1* (reverse): ATG-AAA-ATG-AGC-YAC-WAC-ATA-A [42]. PCR amplification was performed using the Phusion High Fidelity PCR Kit (New England BioLabs, USA). PCRs were performed at a total volume of 50ul consisting of 10ul of 5X Phusion HF Buffer, 1ul 10mM dNTPs, 0.5ul Phusion DNA Polymerase, 2.5ul 10uM forward primer, and 2.5ul 10uM reverse primer. Each reaction contained a total of 100-200 ng of extracted nematode genomic DNA as template. Thermal profile conditions followed Lefoulon and colleagues, and consisted of an initial denaturation at 98C for 30 seconds, followed by 35 cycles of 95C for 30 seconds (denaturation), 51.5C for 45 seconds (annealing), and 72C for 120 seconds (extension) with a final 72C extension for 10 minutes [42]. PCR products (∼970bp amplicon) were purified using ExoSAP-IT PCR Product Cleanup Reagent (Applied Biosystems, USA) and sequenced via classical Sanger sequencing at the University of Minnesota Genomics Center (Minneapolis, Minnesota USA). In parallel, nematode *Cox1* amplicons were sequenced using the Oxford Nanopore Technologies (ONT) MinION sequencing platform. Our intent was twofold, first we wanted to investigate the utility of the newly released (2019) MinION Flongle flow cell for molecular barcoding experiments, especially the depth of nanopore sequencing coverage required to achieve “gold-standard” Sanger sequencing results. Data stemming from this approach would directly support downstream field-based nanopore sequencing in the KCOCA. Second, the single molecule sequencing data generated by the MinION could be leveraged to detect heterogeneous nematode populations that would be difficult to taxonomically characterize using morphology or Sanger sequence data alone.

Nanopore sequencing library construction of the 11 PCR amplicons was performed following the ONT native barcoding genomic DNA protocol using the following kits: EXP-NBD104, and SQK-LSK109. Three sequencing runs were performed using one MinION Flongle vR9.4.1 flow cell per run with a 24 hour runtime, or until no available sequencing pores remained. Raw sequence data produced from the MinION are available via the National Center for Biotechnology Information (NCBI) Sequence Read Archive (BioProject: PRJNA670106). Basecalling of raw data was performed using the Guppy software package (v3.0.6). Samples were demultiplexed for barcoded runs using the Epi2Me software v2020.03.10. NanoPlot v1.13.0 was utilized to provide sequencing summary results and plots of read lengths and quality scores for each sequencing run [43]. FASTQ files generated from basecalling were concatenated (per sample) and randomly sampled (resulting files containing a random sampling of ∼2,000 sequences for each of the 11 samples) using seqtk v1.3 (https://github.com/lh3/seqtk) to determine average coverage needed to obtain a consensus sequence that matched the corresponding Sanger sequence at 100% identity. Sequences between 940–1,000bp were extracted (target *Cox1* amplicon size ∼970bp), and aligned using MAFFT v7.402 [44]. Consensus sequences were generated from randomly sampled files and compared to the corresponding sample’s Sanger sequence using Geneious v2020.1.2.

### Phylogenetic analyses

Sanger sequences were trimmed and manually inspected using Geneious v2020.0.2 (Biomatters, Ltd., New Zealand) and aligned using MAFFT v7.402 [44]. The final alignment was trimmed to 633 bp, the phylogenetically informative length following [42]. Eleven sequences, based on our Sanger sequence data, of *Cox1* from samples (Subfamily: Onchocercinea) collected in South America were included in the final analysis (Fig 4). To root the phylogenetic tree, two species were included as an outgroup: *Breinlia jittapalapongi* and *Brugia malayi*. A maximum likelihood analysis was conducted using the program RAxML v8.2.12 [45] with 1,000 bootstrap iterations and the General Time Reversible + Gamma + Invariable Sites (GTRGAMMAI). Estimates of genetic distance separating clades identified within the resulting phylogeny were calculated using the Kimura-2 parameter (K2P) model of evolution as implemented within the Geneious software package.

**Fig 4.**
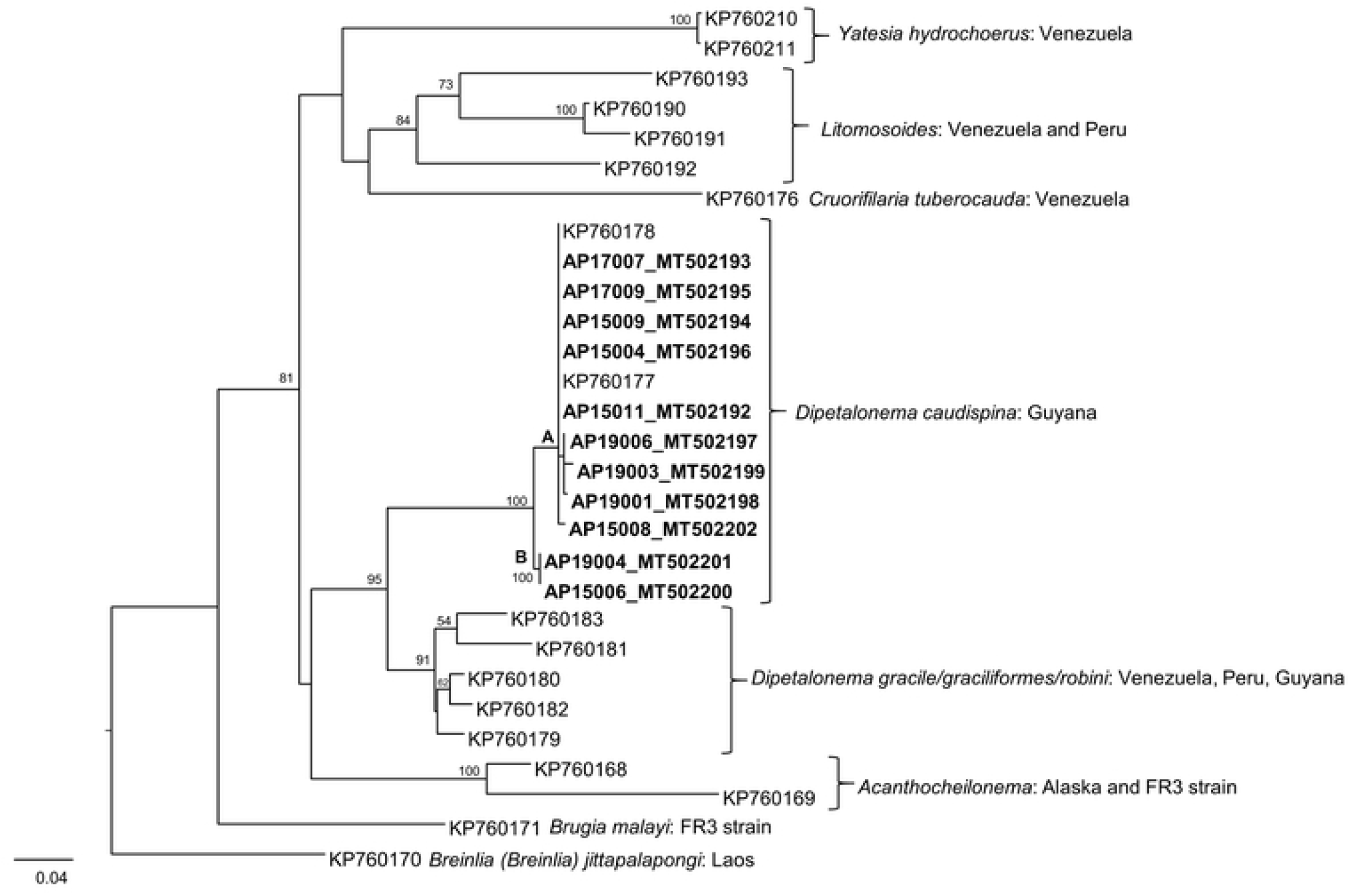
Phylogenetic tree of filarial nematodes constructed from a maximum likelihood analysis using 633 bp of the *Cox1* gene. AP label identifiers were generated for this study (MT labels are assigned Genbank Accession numbers). KP label identifiers were obtained from GenBank and previously reported in [42]. Nodes with bootstrap values ≥ 50 are denoted. *D. caudaspina* is comprised of two clades, A and B, denoted on the tree. *Brenlia jittapalopongi* (KP760170) and *Brugia malayi* (KP760171) are defined as outgroups.

### Ethics Statement

The authors declare no competing interests. Nematode samples were opportunistically collected from animals harvested during normal hunting activities and no animals were collected specifically for this research, thus, this study was determined by the University of Minnesota IACUC committee to be exempt from IACUC approvals. All procedures performed in this study involving human participants were in accordance with the ethical standards of the Institutional Review Boards of the University of Minnesota, Grand Valley State University and with the 1964 Helsinki declaration and its later amendments or comparable ethical standards.

## Results

### Ethnographic results

All interview respondents described the filarids as “*poroto maúre* (Waiwai)” or spider monkey cotton (English) (Fig 1a). They stated that spider monkeys use the cotton to spin into *‘keweyu’* or aprons, a traditional garment worn by Waiwai women (Fig 1b). According to our respondents, the nematodes are not separate from the spider monkey ‘person.’ Instead, respondents indicated that they are an integral part of the monkey’s clothing (or bodily exterior) that hides its innate humanity. Respondents indicated that the cotton was always found with spider monkeys, sometimes found with squirrel monkeys (*Saimiri sciureus*), but rarely found with other primates. They stated that the parasite was never associated with other animals. Many informants were insistent that the cotton was not a “worm” or “parasite,” although others asked if it was indeed a worm. Regardless, informants made a clear distinction between the cotton and other parasites that were observed in the gastrointestinal system of primates (e.g. strongyloids). These other parasites were viewed with revolusion and care was made to properly dispose of the entrails that contained them. In contrast, while some effort was made to wash the spider monkey cotton off of the butchered animal with water, nematodes were often observed being inadvertently cooked and mixed with the meat.

When asked how the animals got the cotton, the following response was typical, “I don’t know, they just have it. It’s a part of them. They must be born with it, and it grows with them, just like their hair.” One informant explained, “For them [spider monkeys], it is their cotton. It is their decoration, their clothes. Just like for us, the apron is our decoration. It doesn’t look the same to us, because we are not monkeys. But to the other monkeys, it is like aprons are to us.”

Other parasites, however, were said to be acquired through the ingestion of certain foods, or transmitted by arthropod vectors. These parasites were viewed as extrinsic to the spider monkey person and perceived as potentially pathogenic to both spider monkeys and humans. Similarly, while respondents suggested that other parasites might be pathogenic, *poroto maúre* was not viewed as disease causing in spider monkeys or zoonotic.

### Histopathology results

All necropsy tissues from spider monkeys harvested in 2015 and 2017 had evidence of filariasis with adult filarid nematodes, microfilariae or both. Adult filarids were found unassociated with any organs and occurred exclusively within the intraperitoneal cavities of the hosts. On histologic evaluation, adult filarids were between 300-800 microns diameter with a smooth eosinophilic cuticle, coelomyarian musculature, small lateral chords, and a lateral internal ridge of the cuticle at the chords. Within the pseudocoelom the intestine was small and there were multiple sections of the reproductive tract, some of which contain microfilariae (Fig 3a). Microfilariae, found within the vasculature of some tissue sections, were 4-6 microns in diameter with tapered ends and numerous internal characteristic basophilic rounded structures (Fig 3b).

**Fig 3.**
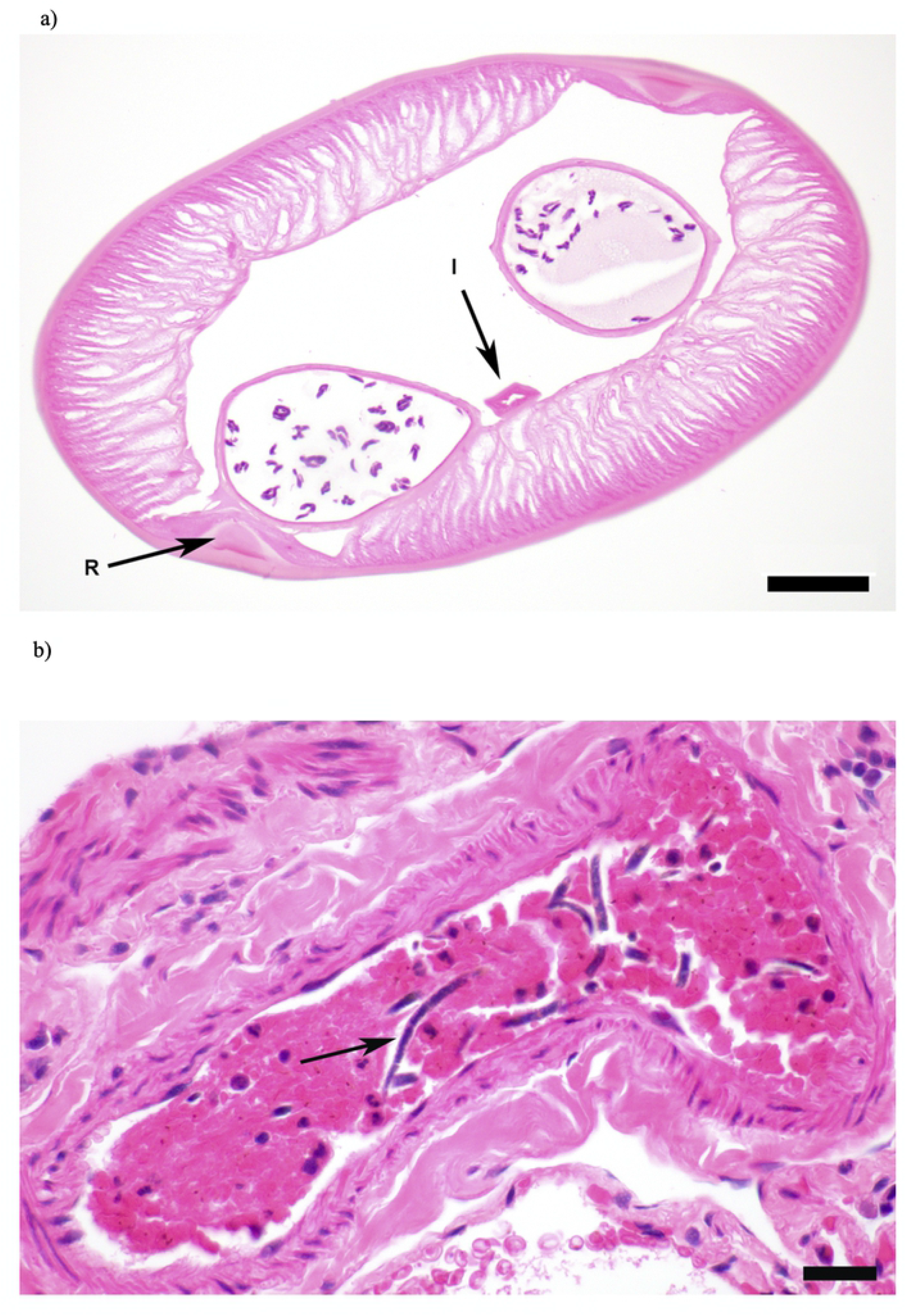
a) Histologic section of an adult nematode from the abdomen of a spider monkey. Characteristic features evident in this image include a smooth cuticle, coelomyarian musculature, lateral internal ridge of the cuticle (R), a small intestine (I) and two uteri with internal microfilariae. Bar = 50 microns. **b) Histologic section of an artery within the lung of a spider monkey**. Within the lumen of the artery, there are numerous microfilariae among red blood cells. The arrow denotes a representative organism. Bar = 20 microns.

### Molecular results

Eleven nematode *Cox1* sequences were generated via Sanger sequencing for phylogenetic analysis and these were deposited to the NCBI GenBank repository (Accession #s: MT502192-MT502202). Sanger sequence data did not exhibit extensive double peaks across our sequence chromatograms, thus indicating a homogeneous nematode population within each spider monkey sample. We performed three MinION Flongle runs of the same 11 *Cox1* amplicons (∼970bp) that were used for Sanger sequencing (S1 Fig). A total of 898,911 reads were generated across the three runs with a mean read quality score of 8.5. We determined that an average of 60X coverage was required to reach at least 99% agreement with the corresponding Sanger sequence across the 11 nematode samples. However, discrepancies consisting of 1 to 2 base insertions/deletions associated with repetitive homopolymer regions (5 to 7 bases in length) were still observed at 60X coverage at approximately 5 regions across individual alignments.

A maximum likelihood analysis was conducted for *Cox1* and generated the tree depicted in Figure 4. The phylogenetic tree depicts a monophyletic relationship for the species *Dipetalonema caudispina* containing 2 distinct, statistically supported clades of Guyanan nematodes (labeled A and B). Approximately 1.5% K2P genetic distance separates *D. caudispina* Clades A and B; whereas *D. caudispina* and its closest sister lineage (the *D. gracile* complex) are 6.9% divergent.

## Discussion

Our study integrated ethnography, histopathology, and molecular phylogenetics to characterize the filarial nematode identified by the Waiwai as spider monkey cotton from both a western phylogenetic perspective and from the ontology of the Waiwai. Morphological and molecular characterization support the taxonomic identification of this filarid as *Dipetalonema caudispina* [46]. We found no evidence of a mixed species nematode population within the individual masses extracted from the spider monkey wild meat samples (Fig 1a). The genus, *Dipetalonema* (Family: Onchocercidae) is composed of six species of filarial parasites known to infect several non-human primate genera throughout the neotropics [42], including: *Ateles, Allouata, Cebus, Saimiri, Saguinus, Lagothrix* and *Aotus* [47]. *Dipetalonema* spp. are widely distributed throughout Central and South America, ranging from Southern Mexico to Paraguay [47]. Hematophagous arthropod vectors in the genus *Culicoides* (Diptera: Ceratopohonidae) are responsible for transmitting *Dipetalonema* spp. between primate hosts; however, the specific vector species involved in transmission and the role the vector plays in the maintenance of *Dipetalonema* populations are poorly understood [48]. Consistent with Waiwai ontological understanding, previous western scientific studies found no evidence of zoonotic potential of spider monkey cotton. Although the family Onchocercidae contains numerous well-characterized human pathogens (e.g. *Wuchereria bancrofti, Brugia malayi, Loa loa*), there is little evidence that any of the six currently recognized members of *Dipetalonema* are zoonotic [49]. Additionally, ethnographic data revealed that the Waiwai perceive other parasites found within spider monkeys as pathologic to both monkeys and humans, specifically, gastrointestinal parasites. Future studies will assess the pathogenic and zoonotic potential of these parasites.

Further, our results contribute to a greater understanding of neotropical primate parasitology. In a review of neotropical primate parasites, Solórzano-García and Pérez-Ponce de León called for the collection of molecular data alongside necropsy data as means to advance current knowledge of neotropical primate parasites [50]. They reported that only 8 of 61 necropsies performed on neotropical primates also included molecular identification of parasites recovered from primate host species [50]. Such efforts are vital, as suitable biological samples are difficult to obtain from free-ranging populations and limited with respect to non-invasive sampling techniques.

### *Dipetalonema* and Waiwai ontology

Our ethnographic results show that the Waiwai do not perceive of *Dipetalonema* as pathogenic to spider monkeys nor potentially transmissible to humans. Instead, they consider the nematode a component of the spider monkey person, much like the monkey’s fur or blood. These beliefs are consistent with perspective multinaturalism, an ontology that has been commonly described for Amazonian societies [27–30], including the Waiwai [36,37]. Multinaturalist societies consider all beings (e.g. jaguar, kapok tree, water spirit) as persons, with a culture and spirit identical to those of humans [28,30]. They have different natures, however (thus multinaturalism), in that they have different bodily exteriors, and these different bodies cause them to have different perspectives. Thus, while spider monkeys consider themselves human, and have societies that operate according to the same norms of the Waiwai, their different bodies make them appear, from the perspective of the Waiwai, as monkeys. As described by Viveiros de Castro, animals in multinaturalist ontology, “experience their own habits and characteristics in the form of culture - they see their food as human food (jaguars see blood as manioc beer, vultures see the maggots in rotting meat as grilled fish, etc.), they see their bodily attributes (fur, feathers, claws, beaks etc.) as body decorations or cultural instruments…[28] (p 470).” In Waiwai ontology, Dipetalonema is a normal bodily attribute of spider monkeys that, from the perspective of the monkey, is cotton that is used to weave an apron, just as the Waiwai would use for their own decoration.

The Waiwai’s understanding of *Dipetalonema* and of host-parasite relations are consistent with our phylogenetic results, allowing us to establish commensurability between scientific and indigenous ontology. Further, by elucidating how the Waiwai conceptualize some parasites as not intrinsic to the spider monkey person, we have identified targets for potential zoonoses. Populations that have existed in environments for hundreds/thousands of years should be expected to have developed cultural beliefs and knowledge that mitigate the potential for zoonotic emergence. While these belief systems may diverge considerably from western metaphysics, understanding divergent ontologies and the role that they may play in promoting sustainable ecological relationships for the societies that hold them can help scientists prevent emerging infectious diseases.

Recently, an increasing number of researchers have sought to utilize local ecological knowledge for managing emerging infectious diseases [17,22,23,51]. For example, Henri and colleagues assessed the extent to which Inuit traditional knowledge could contribute to an understanding of avian cholera outbreaks among Common Eiders (*Somateria mollissima borealis*) [52]. They found that Inuit hunters had extensive knowledge of likely avian cholera outbreaks, several of which were undetected by western biologists. In a study of local ecological knowledge of zoonoses among Maasai pastoralists in Tanzania, Mangesho and colleagues found that while pastoralists generally understood that diseases could be passed between animals and humans, although they had no specific term in their language for zoonotic disease [22]. In addition, the risk of potential zoonoses to humans, particularly diseases that may spread from culturally important practices, was viewed as low and outweighed by the value of the cultural practices for social cohesion. Such research shows the importance of understanding the broader cultural context of local ecological knowledge, particularly how knowledge is situated within a group’s broader ontological framework and cultural practices.

Despite the demonstrated importance of incorporating local ecological knowledge for conservation and environmental management, and its promise in emerging infectious disease research, studies integrating local ecological knowledge with other approaches remain rare in the wild meat literature. In addition, few researchers or theorists have sought to both “take ontology seriously” and assess how commensurability between divergent ontologies might be achieved for managing emerging environmental health challenges [25]. Given the increasing threat of emerging zoonotic disease, and with many zoonotic disease hotspots [13] overlapping with indigenously owned ecosystems, such work has taken on dire significance. By seeking to both understand ontologies that differ from Western metaphysics AND take them seriously as legitimate worldviews, scientists will be better posed to work closely with indigenous groups for preventing zoonotic emergence. As stated by Ludwig and colleagues, “Even if an Indigenous biological ontology diverges from Western science, it still often refers to empirically discovered property clusters that are of outstanding epistemic importance for a community [25](p.46).”

However, environmental and cultural change, both intrinsic (e.g., population increases, technological changes) and extrinsic (e.g., climate change, migration) may impact human-environment interactions in ways that increase the risk of disease emergence. Therefore, integrating multiple knowledge systems and leveraging morphologic and advanced genomic tools to better understand historic human-animal-parasite relationships is critical for preventing zoonotic emergence.

## CONCLUSION

Wild meat is critical to the free security and livelihoods of hundreds of millions of people throughout the world [1,2]. This is particularly true for indigenous groups and other local populations that depend on wild meat for their livelihoods and are on the frontlines of the fight against zoonotic emergence from wild meat. Therefore, it is imperative that scientists work collaboratively with these groups for the co-management of zoonotic risk and the prevention of zoonotic disease emergence. Such collaborations will be most effective when indigenous and scientific ways of knowing are integrated and multiple methodologies are employed. Specifically, co-management partnerships between scientists and indigenous groups will benefit from a recognition of the value of local ecological knowledge alongside scientific knowledge, and, equally importantly, affirmation of indigenous ontologies and understanding the salience of divergent ontologies for the societies that hold them. In addition, such partnerships require the employment of novel scientific approaches to infectious disease screening.

Our results show the value of integrating local ecological knowledge, ontology and molecular characterization for understanding *Dipetalonema* in the KCOCA. We leveraged morphological and molecular phylogenetic approaches to confirm the species identity of the culturally significant spider monkey cotton. Further, our study demonstrates the value of traditional ecological knowledge and indigenous ontologies for understanding zoonotic risk, and has allowed us to identify targets for potential zoonotic pathogens. Our results suggest a commensurability between Waiwai and scientific ontology that will be extremely important in future work of detecting zoonotic agents and mitigating their transmission in this region. We will use this approach (combining genomic tools with traditional ecological knowledge) for long-term management of shared human-animal health in the KCOCA. More broadly, this work highlights the importance of integrating multiple knowledge systems to better understand and prevent emerging zoonotic disease.

## Acknowledgements

We thank the Environmental Protection Agency of Guyana and the Ministry of Amerindian Affairs for granting us permission to conduct this research. We are grateful to Christina Valeri for helping us obtain research permits and Suzanne Stone for helping conduct laboratory work. We are sincerely grateful to the Waiwai of Masakenari Village particularly, Paul Checkema, Dolandea Suse, Charakura Yukuma, Fayu Yukuma, and Romel Shoni, for permitting us to conduct this study and welcoming us into their lives.

## Supporting information

**S1 Table. Genetic divergence values**. Genetic divergence values calculated using the Kimura-2 parameter (K2P) model of evolution.

